# GC content but not nucleosome positioning directly contributes to intron-splicing efficiency in *Paramecium*

**DOI:** 10.1101/2021.08.05.455221

**Authors:** Stefano Gnan, Mélody Matelot, Marion Weiman, Olivier Arnaiz, Frédéric Guérin, Linda Sperling, Mireille Bétermier, Claude Thermes, Chun-Long Chen, Sandra Duharcourt

## Abstract

Eukaryotic genes are interrupted by introns that must be accurately spliced from mRNA precursors. With an average length of 25 nt, the >90,000 introns of *Paramecium tetraurelia* stand among the shortest introns reported in eukaryotes. The mechanisms specifying the correct recognition of these tiny introns remain poorly understood. Splicing can occur co-transcriptionally and it has been proposed that chromatin structure might influence splice site recognition. To investigate the roles of nucleosome positioning in intron recognition, we determined the nucleosome occupancy along the *P. tetraurelia* genome. We showed that *P. tetraurelia* displays a regular nucleosome array with a nucleosome repeat length of ∼151 bp, amongst the smallest periodicities reported. Our analysis revealed that introns are frequently associated with inter-nucleosomal DNA, pointing to an evolutionary constraint to locate introns at the AT-rich nucleosome edge sequences. Using accurate splicing efficiency data from cells depleted for the nonsense-mediated decay effectors, we showed that introns located at the edge of nucleosomes display higher splicing efficiency than those at the centre. However, multiple regression analysis indicated that the GC content, rather than nucleosome positioning, directly contributes to intron splicing efficiency. Our data reveal a complex link between GC content, nucleosome positioning and intron evolution in *Paramecium*.

## INTRODUCTION

In eukaryotes, genomic DNA is compacted by histones into chromatin. The basic unit of chromatin is the nucleosome, which comprises a histone octamer made of the four core histones (H2A, H2B, H3 and H4) and 146-147 bp of DNA wrapped around it (1, 2). Nucleosomes are not randomly located along the genome but positioned with respect to DNA sequence. The affinity of DNA for histone octamers and the energy needed to bend different DNA fragments around the histone octamer are influenced by the primary DNA sequence, which is therefore an important determinant of nucleosome positioning along the genome (3–9). Nucleosome positioning is highly dynamic and its regulation is crucial to control chromatin accessibility and recruitment of chromatin modifiers and transcription factors (10–13). In most genomes, genes display a Nucleosome Free Region (NFR) at the 5’ of their Transcription Start Site (TSS) due to the formation of complexes made by transcription factors around promoter regions (13– 15). A second NFR is present at Transcription Termination Sites (TTS), likely due to the adverse nucleotide composition of the poly(A) signal (16, 17). Nucleosomes are organized in regular arrays with a periodic distance called the nucleosome repeat length (NRL). Such periodicity is especially evident over gene bodies, and it is species and cell-type specific (18, 19). Genome-wide studies have shown that nucleosomes are preferentially positioned in exons compared to introns in diverse organisms including *Schizosaccharomyces pombe, Drosophila*, worms and human (20–25). Several lines of evidence indicated that a well-positioned nucleosome might slow down RNA polymerase II and favour exon inclusion and alternative splicing (26, 27), suggesting a functional role of nucleosome arrays during mRNA maturation. This is in agreement with recent studies showing that intron splicing can occur in a co-transcriptional manner (28, 29). Some studies have suggested that GC richness at exons, and not nucleosome positioning per se, is important for intron splicing (30, 31). Yet, the contribution of nucleosome positioning to intron splicing efficiency has not been investigated thoroughly. The nonsense-mediated decay (NMD) machinery recognizes and degrades transcripts containing premature termination codons (32, 33). Therefore, most of the mis-splicing or un-splicing events are removed rapidly by this powerful surveillance mechanism to avoid the production of erroneous proteins. To date, most studies estimated splicing efficiency from NMD-proficient cells, which eliminate most mis-splicing or un-splicing events, and therefore cannot provide a solid evaluation of intron splicing efficiency.

The ciliate *Paramecium tetraurelia* is a unicellular eukaryotic model organism. Like all ciliates, two distinct types of nuclei co-exist within the same cytoplasm in *P. tetraurelia* (34). The diploid germline micronucleus (MIC) is transcriptionally silent during vegetative growth and transmits the germline genome to sexual progeny through meiosis, while the highly polyploid somatic macronucleus (MAC) is responsible for gene expression (35). The >90,000 introns annotated in the MAC genome are among the shortest reported in eukaryotes (18 to 33 nt, 25 nt on average) (36). How such a large number of tiny introns can be efficiently spliced is not known. In *P. tetraurelia*, no alternative splicing has been reported so far and introns are associated with weak splice signals. A strong counter-selection for introns that cannot be detected by the NMD machinery was previously shown, suggesting that introns rely on NMD to compensate for suboptimal splicing efficiency and accuracy (36, 37). Whether nucleosome positioning or other factors, such as GC content, can regulate splicing efficiency and shape intron evolution in *Paramecium* has not been studied so far.

Here, we investigated a possible role of nucleosome positioning in the recognition of introns in *P. tetraurelia*. We mapped the nucleosome occupancy in the somatic nuclei through paired-end MNase-seq. We found that *P. tetraurelia* displays a regular nucleosome array along genes, with a nucleosome repeat length of ∼151 bp, amongst the smallest periodicities reported in eukaryotes. We compared the positioning of nucleosomes with that of introns and observed that introns are frequently located at the edges of nucleosomes, i.e. associated with linker sequences. Using the accurate splicing efficiency data determined from NMD-depleted cells, we show that introns located at the edge of nucleosomes display higher splicing efficiency than those at the centre. Multiple regression analysis revealed that this difference in splicing efficiency is not explained by the nucleosome positioning per se but rather by the fact that the sequences at the edges of nucleosomes are more AT-rich. This study allowed us to reveal a complex link between nucleosome positioning, GC content and intron splicing efficiency. We propose a model in which a selection constraint during *Paramecium* genome evolution has displaced nucleosome positioning relative to introns, so that the intron sequences are frequently located at the AT-rich edge of nucleosomes to favour efficient splicing.

## MATERIAL AND METHODS

### Paramecium strains, cultivation and autogamy

All experiments were carried out with the entirely homozygous wild type strain 51 of *P. tetraurelia*. Cells were grown at 27°C in wheat grass powder (WGP) infusion medium bacterized the day before use with *Klebsiella pneumoniae* and supplemented with 0.8 mg/mL β-sitosterol (38, 39).

### Macronuclei preparation

Cells were exponentially grown for 12 divisions then cultures at 1,000 cells/mL were filtered through eight layers of sterile gauze. Cells were collected by low-speed centrifugation (550 g for 1 min) and washed once with 10 mM Tris-HCl pH 7.4. The pellet was diluted 3-fold by addition of lysis buffer (0.25 M sucrose, 10 mM MgCl_2_, 10 mM Tris pH 6.8, 0.2% Nonidet P-40) and processed at 4°C as described in (40) with some modifications. Briefly, cells were lysed with 10 strokes of a Dounce homogenizer. Particular care was taken to make sure that macronuclei were still intact under the microscope. Washing buffer (0.25 M sucrose, 10 mM MgCl_2_,10 mM Tris-HCl pH 7.4) was added to a final volume of 10 times the initial pellet. Macronuclei were collected by centrifugation at 2,000 g for 1 min and washed once in washing buffer. The pellet was diluted 2-fold in 2.1 M sucrose, 10 mM MgCl_2_, 10 mM Tris pH 7.4 and loaded on top of a 3-mL sucrose layer (2.1 M sucrose, 10 mM MgCl_2_, 10 mM Tris-HCl pH 7.4) and centrifuged in a swinging rotor for 1 hr at 210,000 g. The macronuclear pellet was washed once, centrifuged at 2,000 g for 1 min and resuspended in washing buffer at 10^7^ nuclei/mL. The macronuclei recovery is quite low, of the order of 10-20%.

### MNase digestion on chromatin isolated from macronuclei

Samples containing 10^5^ macronuclei were incubated in the digestion buffer (0.25 M sucrose, 10 mM MgCl_2_,10 mM Tris pH 7.4, 1 mM CaCl_2_) with increasing amounts (0, 0.5, 1, 2, 5, 7.5, 10 U) of MNase (Sigma) at 30°C for 10 min. Reactions were stopped by the addition of 3 volumes of 0.5 M EDTA pH 9.0, 1% N-laurylsarcosine (Sigma), 1% SDS, 1 mg/mL proteinase K (Merck) and incubated at 55°C overnight. DNA from each sample was gently extracted once with phenol, and dialyzed twice against TE (10 mM Tris-HC1, 1 mM EDTA at pH 8.0) containing 25% ethanol, and once against TE. Samples were then treated with RNase A and DNA was quantified with a Nanodrop spectrophotometer (Thermo Scientific) and separated on a 1.2% agarose gel. The reactions containing mostly mono-nucleosomal DNA fragments (see Fig. 1) were selected and mono-nucleosomal DNA fragments were purified from 3% low melting-temperature agarose gels and treated with β-agarase (Sigma) for sequencing.

**Figure 1.**
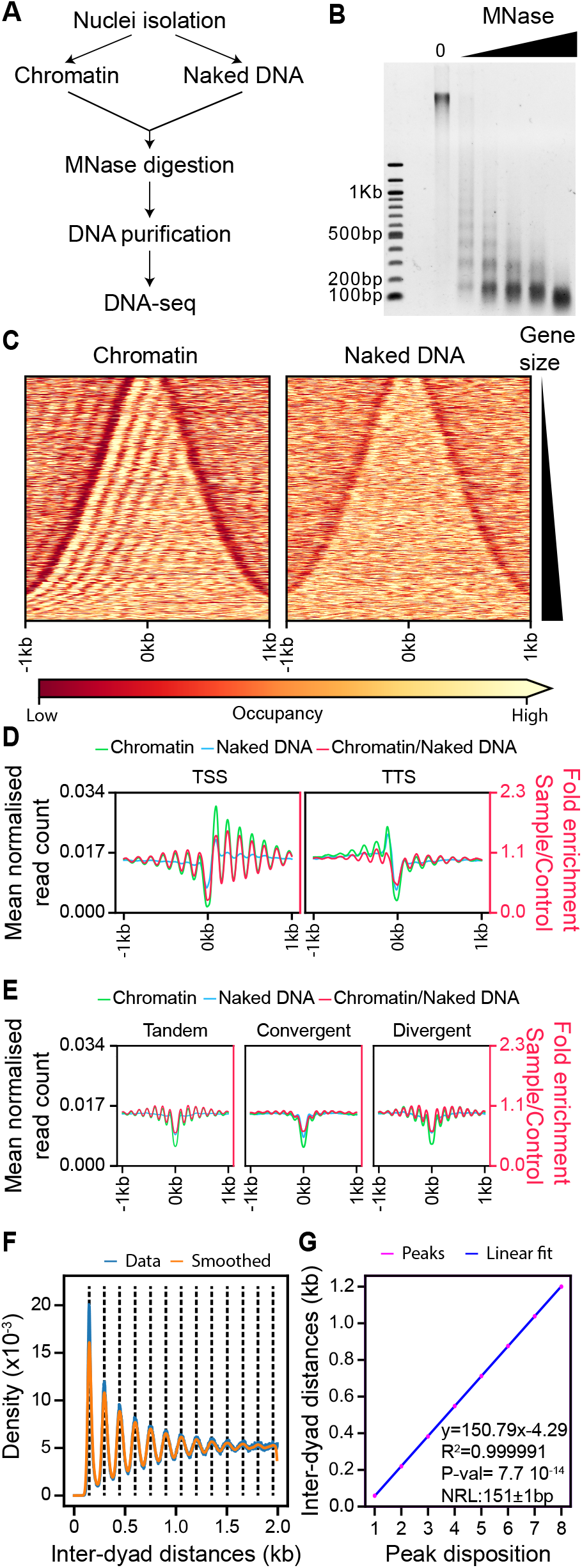
Nucleosome occupancy along the *Paramecium* MAC genome. **(A)** Schematic representation of the MNase-seq experiment. **(B)** MNase digestion of MAC chromatin with increasing MNase enzyme concentration. **(C)** Heatmap showing nucleosome occupancy ± 1 kb around the centre of each gene ordered by gene size (small genes on the top and large genes on the bottom) for 38,256 genes located on scaffolds that are at least 200 kb long. Left panel: average of two chromatin treated samples (Chromatin). Right panel: average of two naked DNA control samples (Naked DNA). **(D)** Average nucleosome occupancies around Transcription Start Sites (TSSs) identified by 5’ CAP-seq on the left, and Transcription Termination Sites (TTSs) identified by poly(A) detection on the right: in green, the average profile of chromatin treated sample (Chromatin); in blue, average profile of naked DNA treated sample (Naked DNA); and in red the Chromatin/Naked DNA ratio, enrichment of which is shown on the second axis on the right (red axis). **(E)** Average nucleosome occupancies ±1 kb around the centre of intergenic regions: same colour code as in panel D. Intragenic regions have been divided into three groups based on the relative positions of gene pairs: tandem (left), convergent (middle) or divergent (right). **(F)** Inter-centre distance between well-positioned nucleosomes (Materials and Methods) on the same scaffold. In blue, distance distributions from actual data (from 1 bp to 2 kb, binning=1); and in orange, the gaussian smoothed signal. Black dashed lines indicate the local maxima (peak centres) of the smoothed data (Materials and Methods). (**G**) In pink, the first 8 local maxima from Fig. 1 F ordered by increasing distance, and in blue the linear fitted model. At the bottom right, information about linear fitting and estimated NRL (Mean±SD) is given. P-value is calculated using a two-sided Z-test.

### MNase digestion on naked DNA

Following purification on a sucrose layer, the macronuclear pellet was washed once, centrifuged at 2,000 g for 1 min, and was resuspended in three volumes of lysis solution (0.5 M EDTA at pH 9.0, 1% SDS, 1% N-laurylsarcosine (Sigma), 1 mg/mL of proteinase K (Merck) then incubated at 55°C overnight. DNA was gently extracted with phenol, and dialyzed twice against TE (10 mM Tris-HC1, 1 mM EDTA at pH 8.0) containing 20% ethanol, and once against Tris 10mM pH 8.0. 1.6 μg of DNA was digested with increasing amounts of MNase (0 to 1×10^−3^ U) in the digestion buffer at 30°C for 10 min. The reactions were stopped with 250 mM EDTA. The samples were analysed on a 1.2 % agarose gel and reactions containing fragments of 100-200 bp were gel-purified for DNA sequencing (see Fig. S1).

### MNase library preparation and sequencing

Sequencing libraries were generated using the sequencing kit: TruSeq SBS Kit v5 – GA (36 Cycle) (FC- 104-5001, Illumina). Samples were then sequenced on an Illumina GA-IIx sequencer using paired-end (PE) 74 bp setting. Alignment was performed using ELAND and mapping to the MAC genome of strain 51 v1.0 (ptetraurelia_mac_51.fa), available at ParameciumDB (https://paramecium.i2bc.paris-saclay.fr/) (41).

### Nucleosome positioning calling

After aligning reads to the reference MAC genome, PCR duplicates with the same start and end positions were removed. Only reads from mono-nucleosomes were kept, therefore, read pairs longer than 150 bp and shorter than 75 were excluded. We used only the data within the scaffolds larger than 200 kb. A nucleosome score was calculated using the central 75 bp of each read pair. Signal was then smoothed with a gaussian filter and a sigma of 30 over 90 bp for visual assessment of nucleosome position calling. Local Maxima and local Minima were identified by convoluting the nucleosome score with a first derivative of a gaussian (sigma 30 over ±90 bp). The points of inflection were identified by convoluting the nucleosome score with a kernel containing the second derivative of a gaussian (sigma 30 over ± 90 bp). Peaks were called as a local maximum between two inflection points with opposite inclination. Peaks were called independently in the two chromatin samples, and then a list of well-positioned nucleosomes was compiled using those nucleosomes whose dyad (i.e. centre) differs by less than 10 bp between the two biological replicates (80% of all nucleosomes). These well-positioned nucleosomes were used for the downstream analyses.

### Computation of Nucleosome Repeat Length

To compute the Nucleosome Repeat Length (NRL), we first calculated the distance of each nucleosome to all the other nucleosomes on the same scaffold, then used the distances obtained to generate the density distribution. This density distribution was then smoothed using a gaussian filter (sigma=10 over ±30bp) and local maxima identified convoluting the density distribution with the first derivative of a gaussian (sigma=10 over ±30bp). The first 8 local maxima were then ordered by increasing distances and fitted using a linear model. The slope of the fitted model corresponds to the NRL.

### Nucleosome distribution calculation

Gene annotation v2.0 of MAC was from (41), and the transcription start sites and transcription termination sites were from (42). To compare with the distribution of real exon sizes, a set of simulated exons was created assuming uniform exon sizes within each gene, i.e. for a given gene with *n* exons, we divided its total exon length by *n* to get the length of *n* simulated exons of the corresponding gene. The NMD data were obtained from (37), splicing efficiency of each intron was calculated as the Splicing events / Total number of observations (i.e. spliced + unspliced reads). Mean profiles and heatmaps were drawn using a customized script and plotting using matplotlib. All statistical analyses were performed with Python (version 3.7.4).

### Multilinear regression

Starting parameters used for the multiple linear regression can be found in Table S1. Parameters were transformed using appropriate functions in order to maximise their linearity with the intron splicing efficiency, e.g. log transformation of expression levels. Values were then standardized. A randomly selected set of introns (10% of all introns) was kept from the multilinear regression model fitting, and used as a test dataset to evaluate the model performance. Parameters were tested in multiple combinations. For each combination, after each fitting, we performed a two-sided Z-test per each coefficient with H_0_: C=0 and H_1_: C≠0. Statistically significant coefficients were then retained and the linear model was trained again with the associated parameters. This step was repeated until the number of variables stabilised. Using the intron test dataset, we calculated the Pearson correlation between real and predicted data, and the best predicting model was kept. Estimation of the contribution of each parameter is calculated as in (43), which is based on the absolute value of the product of each coefficient and the Pearson correlation value of its parameter with the splicing. Contributions were then converted to percentages.

## RESULTS

### Genome-wide nucleosome position profiling along the *Paramecium* somatic genome

Using MNase-seq, we derived a first nucleosome positioning profile of the macronuclear (MAC) genome of *P. tetraurelia* during vegetative growth. Both chromatin samples and naked MAC DNA controls were digested to mono-nucleosome size (∼150 bp, Fig.1 A & B and Fig. S1 A). The results obtained from two biological replicates were highly reproducible (Pearson R = 0.92, Fig. S1 B). We therefore combined data from both biological replicates for downstream analyses. All the data presented in the main figures were obtained with the average of two chromatin samples and two naked DNA controls, respectively. The results of each individual sample are reported in the supplementary figures. Using the gene annotation, together with the Transcription Start Sites (TSSs) identified by 5’ CAP-seq and Transcription Termination Sites (TTSs) identified by poly(A) detection (42), we investigated nucleosome occupancy along transcription units and around their extremities. As described in other eukaryotes, *P. tetraurelia* presents an enriched nucleosome density over the transcription units compared to the flanking regions, showing regular arrays of nucleosomes over transcription units (Fig. 1 C-D and Fig. S1 C-D). As expected, we were able to identify Nucleosome Free Regions (NFRs) upstream of the TSSs of *Paramecium* genes, followed by an array of well-positioned nucleosomes (Fig. 1 C-D and Fig. S1 C). The analysis of TTSs shows regions with very low nucleosome occupancy downstream of the TTSs and a weakly organised array towards the gene body (Fig. 1 D and Fig. S1 D). We further separated gene pairs into 3 groups based on their disposition: tandem (n=20,298), convergent (n=8,900) and divergent (n=8,890) (Fig. 1 E and Fig. S1 E-G). Interestingly, we found that nucleosome arrays are clearly visible upstream of the TTS only when genes are positioned in tandem or divergent pairs (Fig. 1 E and Fig. S1 F), but not in convergent pairs (Fig. 1 E and Fig. S1 G). This observation suggests that the nucleosome positioning at TTS observed for tandem genes might be due to the downstream TSS, as suggested for *Saccharomyces cerevisiae* (44). Alternatively, convergent genes might be influenced by the transcription readthrough of the gene in the opposite orientation.

Based on our nucleosome position calling and using only well-positioned nucleosomes identified in both replicates (see Materials and Methods), we calculated the nucleosome repeat length (NRL) (Materials and Methods). We found that *P. tetraurelia* displays one of the smallest NRL reported in eukaryotes (150.79±0.56 bp on average, Fig. 1 F-G and Fig. S1 H-I), close to the 156±2 bp of *S. pombe* (45), which is much smaller than the 165 bp of *S. cerevisiae* (46). In human, the NRL within gene bodies is smaller than outside (47). We performed a similar analysis sub-setting nucleosomes based on whether their centres overlap with gene bodies or not. We found a negligible difference between the NRL within gene bodies (150.79±0.70 bp, more than 80% of the analysed sequences) and those outside of genes (150.99±1.17 bp) (Fig. S1 J).

### The tiny introns of *Paramecium* genes are frequently associated with inter-nucleosomal DNA

We then analysed nucleosome positioning over gene bodies. In *P. tetraurelia*, exons range from several nucleotides to a few kilobases (Fig. 2 A, Fig. S2 A) and are interspersed with tiny introns, spanning between 18 and 35 bp with a median size of 25 bp (Fig. 2 B). By visual inspection of the nucleosome occupancy profiles, we noticed a tendency of the MNase signal to be stronger over exons leaving the introns preferentially between two nucleosome peaks (Fig. 2 C, Fig. S2 B). This was especially visible when we examined the nucleosome density over introns sorted by the distance of each intron centre to the closest nucleosome centre (Fig. 2 D, Fig. S2 D). This distance is significant higher than what we would expect by calculating the distance of random positions inside gene bodies to the closest nucleosome centre (*p* value < 10^−10^ calculated with Mann-Whitney U test, one sided, alternative H_1_: Introns distance from the closest nucleosome is higher than random chance. Fig. 2 E). Using this distance, we grouped introns into 3 categories: central, proximal and distal introns (as illustrated in Fig. 2 F). We calculated their distribution and compared it with that of exons smaller than 300 bp (roughly the same sample size) categorized in the same way (Fig. 2 G). Introns were found enriched at distal positions, i.e. located in the regions between two neighbour nucleosomes, compared to exons (46% vs 22%, respectively). In contrast, exons were more enriched in central positions compared to introns (47% vs 27%, respectively). These distributions are statistically significantly different: *p* value < 10^−10^ calculated with a χ^2^ test (Fig. 2 G). Moreover, *P. tetraurelia* exons seem to favour mono-nucleosome length sizes with 35% of exons having sizes comprised between 100 bp and 200 bp. Such a size distribution is significantly shorter than what one would expect if exon sizes were simply uniformly distributed within each transcript (see Materials and Methods), in which case only 24% of the exons would fall in this range (*p*<10^−10^, Mann-Whitney U test, one sided, alternative H_1_: simulated exons are bigger than real exons) (Fig. 2 A). Similar results were obtained using only exons of transcription units whose extremities are identified by both 5’ CAP-seq and poly(A) detection (Fig. S2 A). This distribution of exon sizes might reflect some selective constraint to keep introns in phase in distal position, i.e. at the edge of the nucleosome.

**Figure 2.**
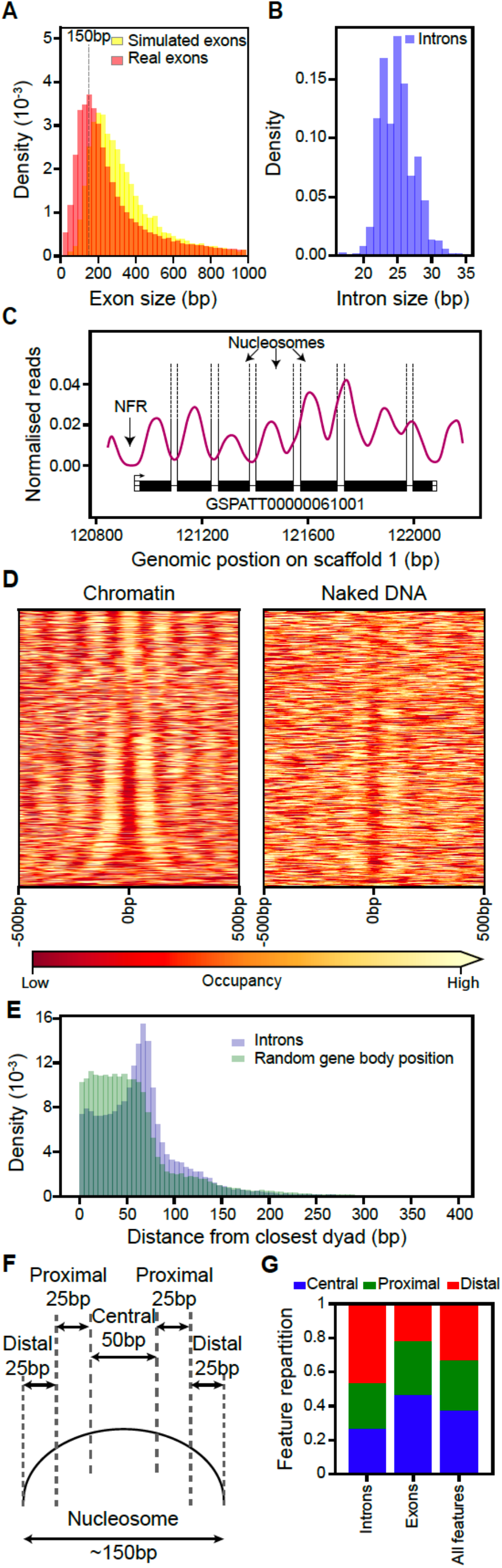
Inter-nucleosomal DNA is frequently associated with intron position. **(A)** Histogram showing exon size distribution (bin size = 25 bp): in red, real exons; and in yellow, simulated exons created assuming uniform exon sizes within each gene (see Materials and Methods). **(B)** Histogram showing intron size distribution (bin size = 1 bp). **(C)** Example track reporting nucleosome occupancy over genes with intron locations indicated by dashed vertical lines. We can observe nucleosome free regions (NFRs) around the gene promoters and introns frequently associated with inter-nucleosomal DNA. **(D)** Heatmap showing nucleosome occupancy ± 500 bp around intron centres. Introns are ordered based on increasing distances from their centre to the closest nucleosome centre. The average of the chromatin samples is shown on the left and the average of the naked DNA samples on the right, with the same colour code as in Fig. 1 C. **(E)** Histogram reporting the distance of an intron centre to the closest nucleosome centre (blue). For each intron, a random position inside the corresponding gene body was selected and the distance to its closest nucleosome centre is reported (green). Bins size= 5 bp. **(F)** Schematic representation of the criteria to assign features for each intron (or exon) into one of the 3 classes, based on the distance (*d*) between its centre and the closest nucleosome centre position: central, *d* ≤ 25 bp; proximal, 25 bp < *d* < 50 bp; distal 50 bp ≤ *d* ≤ 75 bp. **(G)** Relative distribution of introns, exons and both features over categories defined in (F) for the introns overlapping with a fixed nucleosome (about 70% of all introns, see Materials and Methods) and exons with a size below 300 bp overlapping with fixed peaks. See Fig. S2 C including the features with *d* > 75 bp.

### Higher splicing efficiency for introns at the edges of nucleosomes

Previous studies have described the effect of nucleosome positioning on mRNA maturation in multiple organisms (20–25). To address whether nucleosome positioning affects intron splicing in *P. tetraurelia*, we examined the relationship between nucleosome positioning and intron splicing efficiency, using published datasets from both wild-type (WT) and NMD-depleted cells, which provide a measurement of the splicing efficiency of *P. tetraurelia* introns (37). Since NMD has been shown to play an important role in removing mis-spliced isoforms and different evolutionary constraints have been observed for NMD-visible (presence of a premature stop codon after retention) and NMD-invisible (absence of a premature stop codon after retention) introns (37), we further divided our 3 positional categories (central, proximal, distal) of introns into NMD-visible or NMD-invisible groups (Fig. 3 A).

**Figure 3.**
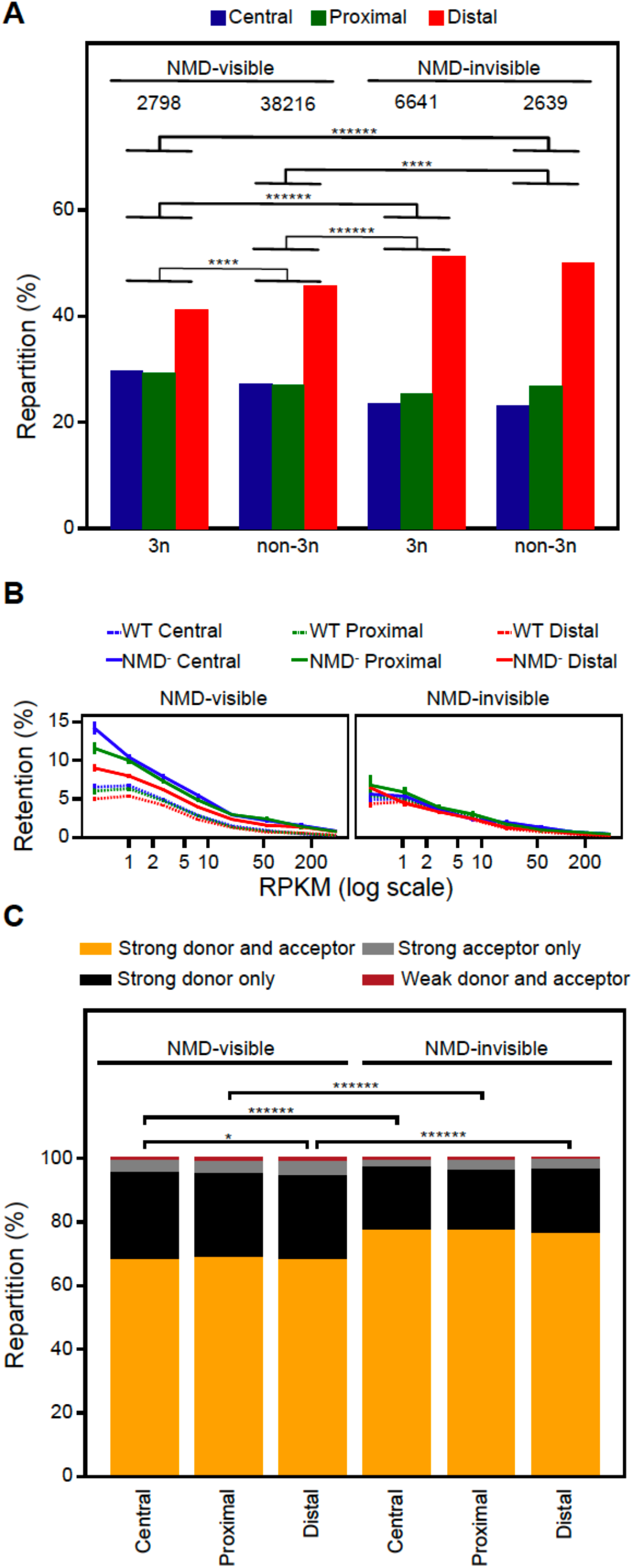
Nucleosome positioning is associated with intron-splicing efficiency. **(A)** Relative distribution of different classes of introns. Introns are grouped based on their length (3n or non-3n) and whether their retention causes a premature stop codon making them visible to the nonsense-mediated decay mechanism (NMD-visible) or not (NMD-invisible). Within each group, introns are classified based on the distance to the closest nucleosome as in Fig. 2 D. P-values are calculated using the χ^2^ test and only the significant ones are indicated. **(B)** The retention rate of introns in WT (dashed lines) and in NMD-depleted (solid lines) cells as a function of gene expression levels. Introns are classified based on their distance to the closest nucleosome as in Fig. 2 D and on whether they are visible to NMD or not. Error bars represent the standard error of the mean. P-values calculated using Mann-Whitley U test, and adjusted with false discovery rate, are displayed in Fig. S3 A. **(C)** Relative characterization of introns with strong splicing acceptors and/or donors, with the same categories as in Fig. 3 B. P-values are calculated using the χ^2^ test. (P-value *<0.05, **< 10^−2^, ***< 10^−3^, ****< 10^−4^, *****< 10^−5^, ******< 10^−6^).

First, we observed that the proportion of distal introns is higher in NMD-invisible introns compared to NMD-visible ones, independent of the introduction of a frameshift (3n versus non-3n introns) (Fig. 3 A). We could not observe statistically significant differences between 3n and non-3n NMD-invisible intron distributions (*p*=0.25, χ^2^ test), and only a minor significant increase of distal introns at the expense of central introns and proximal introns can be detected between 3n and non-3n NMD-visible introns (*p*<10^− 6^, χ^2^ test) (Fig. 3 A). Since no major differences in the intron distribution between 3n and non-3n introns were observed, we decided to consider only the NMD state for subsequent analysis.

As shown in (37), the intron retention rate is inversely correlated with the gene expression level and is higher for introns that can be detected by the NMD machinery than for those that cannot. In WT cells, both NMD-visible and NMD-invisible introns showed similar retention rates, with higher retention rates for genes with lower expression levels (Fig. 3 B). The retention rate of NMD-visible introns increased significantly upon NMD depletion, while it did not in NMD-invisible introns (Fig. 3 B). We extended this analysis to our intron positional categories. As expected, NMD-invisible introns showed similar splicing efficiency for all intron classes in both WT and NMD-depleted cells (Fig. 3 B right panel and Fig. S3 A). Interestingly, we found that the retention rate of NMD-visible introns decreased while the distance of the intron to the closest nucleosome centre increased (Ret_Central_>Ret_Proximal_>Ret_Distal_, Fig. 3 B left and Fig. S3 A), indicating that NMD-visible introns located at the edges of nucleosomes are more efficiently spliced. This can already be observed in a WT background, while in NMD-depleted cells, where nonsense mRNAs are no longer degraded, this difference is much stronger (Fig. 3 B left panel and Fig. S3 A). For the lowest-expressed genes, the retention rate of central introns is 57.6% higher than that of distal introns, and it drops to 38.8% and 13.9% for the median and highly-expressed genes, respectively (Fig. 3 B left panel and Fig. S3 A).

It has been shown that the splicing efficiency of *P. tetraurelia* introns depends on the sequences at the donor and acceptor sites (36). We thus assessed whether this difference in splicing efficiency between our nucleosome-positional classes could be explained by a different distribution of strong donor (5’ GTA) and/or strong acceptor (3’ TAG) sites (Fig. S3 B) within different intron groups. As expected, NMD-invisible introns were more frequently associated with both strong donors and acceptors whatever the distance of the intron to the closest nucleosome centre (Fig. 3 C). In contrast, we found a minor increase in the association of distal introns with “weak donor and acceptor” and “strong acceptor only” intron groups compared to central introns, for the NMD-visible introns (Fig. 3 C). This slight increase was not associated with a higher retention rate for distal introns compared to central introns. Instead, we did observe a reduced retention rate in distal introns (Fig. 3 B). We conclude that the reduced retention rate in distal introns is not due to a difference of donor/acceptor signals in this class.

### GC content related to nucleosome positioning contributes to intron-splicing efficiency at the edges of nucleosomes

It is well known that nucleosome positioning is highly associated with GC content: nucleosome centres show higher GC than distal regions (3, 6–9). In *P. tetraurelia*, we observed that NMD-visible introns have a higher GC content than their NMD-invisible counterparts (Fig. 4 A). Moreover, as we would expect, the central introns have the highest GC content followed by proximal and distal introns (Fig. 4 A). We therefore analysed the impact of GC content on intron retention rates. We found a direct correlation between GC percentage and retention rate in NMD-visible introns, yet no statistically significant difference could be observed between different intron groups (Fig. 4 B and Fig. S4 A). This suggests that GC-content anti-correlates with intron splicing efficiency. To further evaluate how different parameters, such as GC content, gene transcription level, and nucleosome positioning (Table S1) affect intron splicing efficiency, we iteratively trained a multivariate regression model as previously described (43), by using different combinations of parameters and retaining only those that were statistically significant (Materials and Methods). The best fitting model (R=0.6), which explains 35% of variation in intron splicing efficiency was then used to estimate the contribution of each parameter (full list of parameters in Table S1).

**Figure 4.**
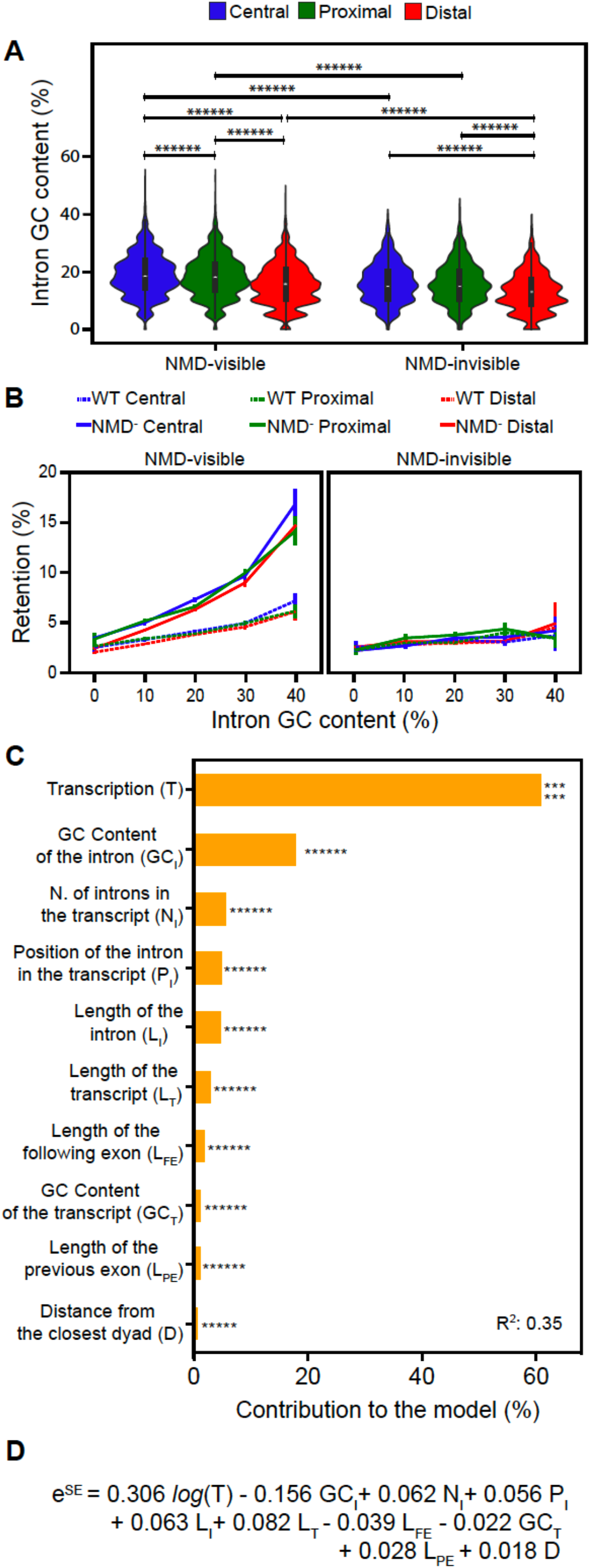
GC content related to nucleosome positioning contributes to intron-splicing efficiency. **(A)** GC content (%) distribution of introns based on the distance to the closest nucleosome centre and NMD visibility. P-values were calculated using the Mann–Whitney U test and adjusted using the false discovery rate. **(B)** The retention rate of introns in WT and NMD-depleted cells as a function of their GC content (excluded GT and AG dinucleotides at both extremities). Introns are classified based on their distance to the closest nucleosome centre and on whether they are visible or not to NMD. Binning = 10%. Error bars represent the standard error of the mean. P-values calculated using the Mann–Whitney U test, and adjusted using the false discovery rate, are displayed in Fig. S4 A. **(C)** Modelling Splicing Efficient (SE) in NMD-depleted cells, showing the parameters used to fit our model, with the contribution of each parameter to the model and their statistical significance (P-Val). **(D)** The full fitted model in explaining intron splicing efficiency, indicating whether each parameter is positively or negatively correlated with splicing efficiency. The parameters are as indicated in (C). (P-value *<0.05, **< 10^−2^, ***< 10^−3^, ****< 10^−4^, *****< 10^−5^, ******< 10^−6^).

The highest contribution came from the level of gene expression that accounts for about 60% of the model. The GC content of the intron accounted for almost 18%. The length of the intron, its position in the transcript and the total number of introns in a transcript accounted for about 5% each in the model, followed by the size of the following exons (∼2%), the GC content of the transcript (∼1%) and the size of the previous exon (∼1%). Remarkably, the only parameter relative to the nucleosome positioning retained by the model was the distance to the closest nucleosome centre that accounts for only 0.64% (Fig. 4 C-D and Table S1). In addition, once we divided the introns into NMD-visible and NMD-invisible, the distance to the closest nucleosome centre was relevant only for the NMD-visible ones where it accounts for 0.67% of the model (Fig. S4 B-C). We therefore conclude that, the GC content, which is tightly linked to nucleosome positioning, contributes to intron-splicing efficiency.

## DISCUSSION

We have performed the first nucleosome position profiling in the *P. tetraurelia* MAC genome during vegetative growth. Despite its high AT richness (>75% AT), the *P. tetraurelia* MAC genome displays a very regular nucleosome positioning pattern as observed in other eukaryotes: NFRs at the TSSs and TTSs, and a regular nucleosome array along genes. Unlike *Tetrahymena thermophila*, another AT-rich ciliate (78% AT) (with an NRL of 199 bp) (48), the NRL in the *P. tetraurelia* MAC genome presents a smaller periodicity (151±1 bp), very similar to that of *S. pombe* (156 bp) (45) and of *Plasmodium falciparum* (155 bp) (>80%AT) (49, 50). This short NRL means that the naked “linker” DNA between nucleosomes in *Paramecium* is extremely small (only a few bp) compared to that of most other eukaryotic genomes, which is at least tens of bp or even larger (51). A negative correlation between the H1/core-histone ratio and NRL has been previously reported (52–54). Importantly, for the three eukaryotes with the smallest NRL, *P. tetraurelia, P. falciparum* and *S. pombe*, no orthologue of histone H1 has been identified so far. This strongly suggests that the absence of H1 might contribute to the extremely short NRL observed in *Paramecium* chromatin organization in the somatic MAC genome.

In yeast and human, actively transcribed genes tend to have shorter NRL than transcriptionally inactive genes, partially due to the binding of H1 generating inaccessible chromatin at inactive genes (47, 55, 56). With the separation of the germline MIC and the somatic MAC genomes in two distinct nuclei, the *Paramecium* MAC genome is very compact. Indeed, >80% of the MAC is covered by annotated genes and 65% of the coding genes are expressed (covered by at least 1 RPKM) during vegetative growth (34, 42), which might explain the extremely short length and narrow distribution of NRL. A significant difference in the nucleosome organization between MAC and MIC genomes has been reported for *T. thermophila* (57). How nucleosomes are organized in the *Paramecium* MIC genome is unknown. At each sexual cycle of *Paramecium*, the parental MAC is destroyed and the new MIC and MAC are generated from the parental germline MIC (35). During new MAC development, an estimated ∼25% of the germline DNA is eliminated during massive genome rearrangements (58). A large amount of extremely short (26 to ∼1000 bp) non-coding germline sequences, called IESs (Internal Eliminated Sequences), need to be precisely excised to correctly assembly functional genes in the new MAC genome of *Paramecium* species (59). How nucleosome positioning is organized in the germline MIC genome relative to IESs and whether nucleosome positioning and/or GC content might play a role in IES excision are open questions (60, 61).

In multicellular eukaryotes, long introns are recognized through exon definition and nucleosomes positioned along exons might contribute to the exon-intron architecture, possibly pointing to a function in exon definition (20–25). By contrast, short introns are recognized through intron definition. With an average length of 25 nt, introns of *P. tetraurelia* are among the shortest reported in eukaryotes (36). The large number of introns (>90,000) are associated with weak splicing signals. In the current study, we examined the role of nucleosome positioning in intron splicing. We found a regular nucleosome array associated with intron positions within genes, with exons wrapped around nucleosomes and introns frequently located at the edge of nucleosomes. By using the accurate splicing efficiency data obtained from NMD-depleted cells (37), we performed a thorough investigation on the effect of nucleosome positioning on splicing efficiency. We showed that the NMD-visible introns located at the edge of nucleosomes display higher splicing efficiency than those at the nucleosome centres. However, we found that this higher splicing efficiency is due to the fact that the introns located at the edges of nucleosomes display lower GC content. Our multiple regression analysis indicated that the nucleosome positioning did not show any contribution to the intron splicing efficiency (Fig. 4 C). Our results strongly support that the GC content, rather than the nucleosome positioning, directly influences intron splicing efficiency in *Paramecium*, which may pave the way to future mechanistic studies to decipher how GC content may impinge on intron splicing efficiency. Whether the effect of GC content and nucleosome positioning on intron splicing efficiency observed in *Paramecium* can be extended to other eukaryotes remains an open question.

Interestingly, we also observed that during evolution, nucleosome positioning has been displaced relative to introns, so that the AT-rich intron sequences are frequently located at the edge of nucleosomes (Fig. 4 A). Although both NMD-visible and NMD-invisible introns present a higher proportion of distal positions, NMD-invisible introns show a significantly higher proportion (51% for 3n and 49% for non-3n introns) than NMD-visible introns (41% for 3n and 46% for non-3n introns) (Fig. 3 A). This strongly suggests that the NMD-invisible introns not located at the AT-rich nucleosome edges, whose retention in transcripts cannot be cleaned up by the NMD pathway, are counter-selected during evolution. Whether introns in *Paramecium* might play a functional role is still unclear. These introns do not seem to contribute to alternative splicing to generate protein diversity or to encode ncRNAs as in other eukaryotic genomes (62–64). Due to their extremely small sizes, these introns are unlikely to play a role in regulating the transcription rate as suggested in recent publications (65–67). How such a large number of tiny introns in *Paramecium* is maintained during evolution and how these introns can be efficiently spliced need to be further investigated.

## DATA AVAILABILITY

The gene annotations and transcription data are available at ParameciumDB (https://paramecium.i2bc.paris-saclay.fr/). The customized script and Jupiter notebooks used for this study are available on the GitHub page (https://github.com/CL-CHEN-Lab/Nucleosome).

## ACCESSION NUMBERS

MNase-seq datasets are deposited in the ENA under the Project Accession PRJEB39679 (68).

## AUTHOR CONTRIBUTIONS

LS, MB, CT, CLC and SD conceived and planned the study. MM, FG and SD conducted the experiments. SG, MW, OA and CLC performed the bioinformatics analyses. CT and CC supervised the bioinformatics analyses. SG, CLC and SD wrote the manuscript, and all the authors reviewed it.

## ACKNOWLEDGEMENTS

The authors would like to thank Laurent Duret for useful suggestions and discussion, Laurent Duret and Eric Meyer for sharing with us the NMD data, and to acknowledge the high-throughput sequencing facility of I2BC for its sequencing and bioinformatics expertise.

