## Supplementary Data for "GC content but not nucleosome positioning directly contributes to intron-splicing efficiency in *Paramecium*"

| Table S1 |
|---|
| Average distance between the intron of interest centres the two closest nucleosomes |
| Distance between the intron of interest centre and the centre of the closest nucleosome |
| GC content of the intron of interest |
| GC content of the following exon |
| GC content of the previous exon |
| GC content of the transcript |
| Intron position in the transcript |
| Length of the following exon |
| Length of the intron |
| Length of the previous exon |
| Length of the transcript |
| Median MNase signal over the intron |
| Median MNase signal over the following exon |
| Median MNase signal over the previous exon |
| Number of introns in a transcript |
| Transcription level |

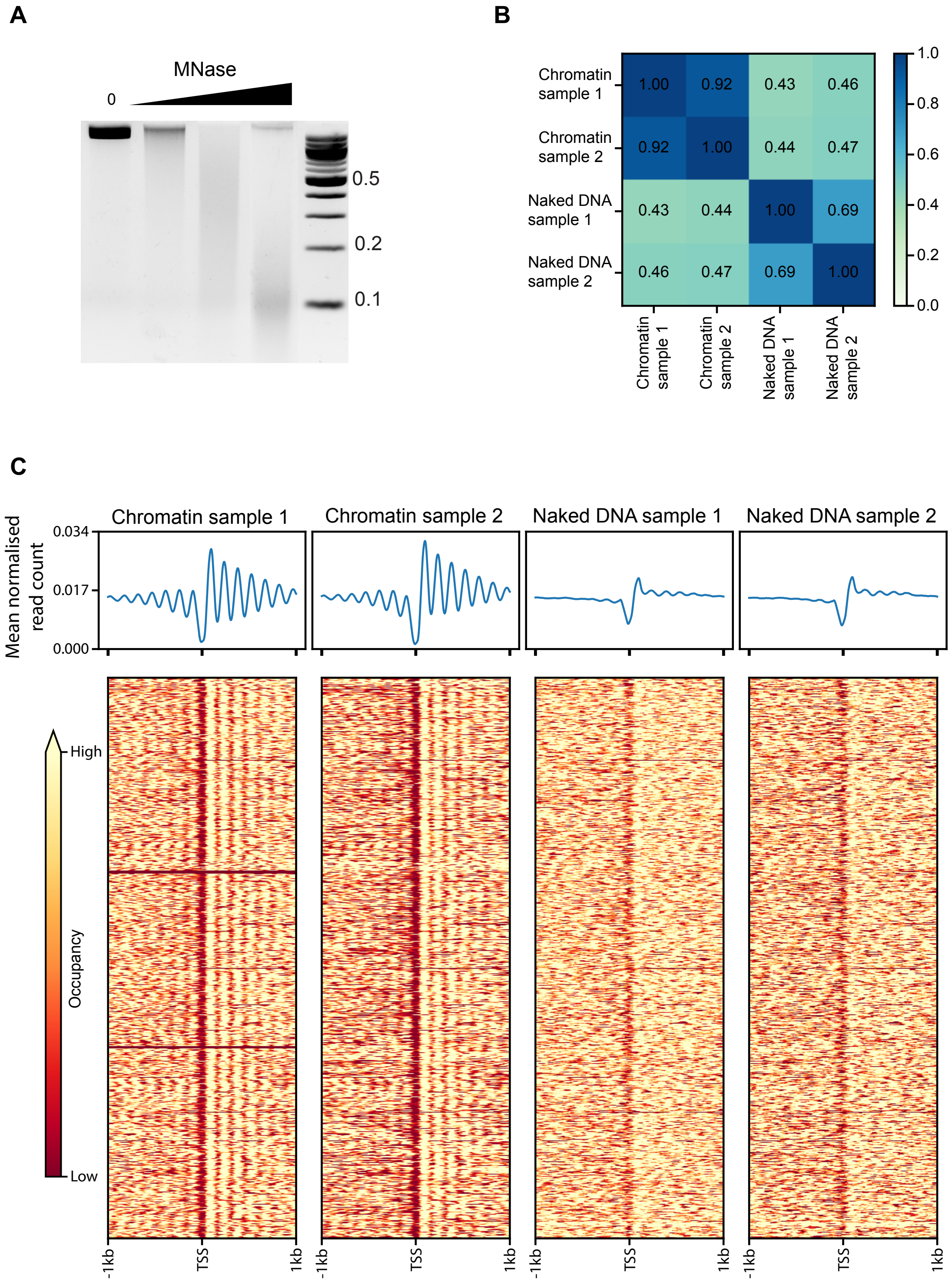

A
MNase
0
0.5
0.2
0.1
B
Chromatin sample 1
Chromatin sample 2
Naked DNA sample 1
Naked DNA sample 2
1.00 0.92 0.43 0.46
0.92 1.00 0.44 0.47
0.43 0.44 1.00 0.69
0.46 0.47 0.69 1.00
1.0
0.8
0.6
0.4
0.2
0.0
C
Chromatin sample 1
Chromatin sample 2
Naked DNA sample 1
Naked DNA sample 2
Mean normalised read count
0.034
0.017
0.000
High
Occupancy
Low
-1kb
TSS
1kb

**D**

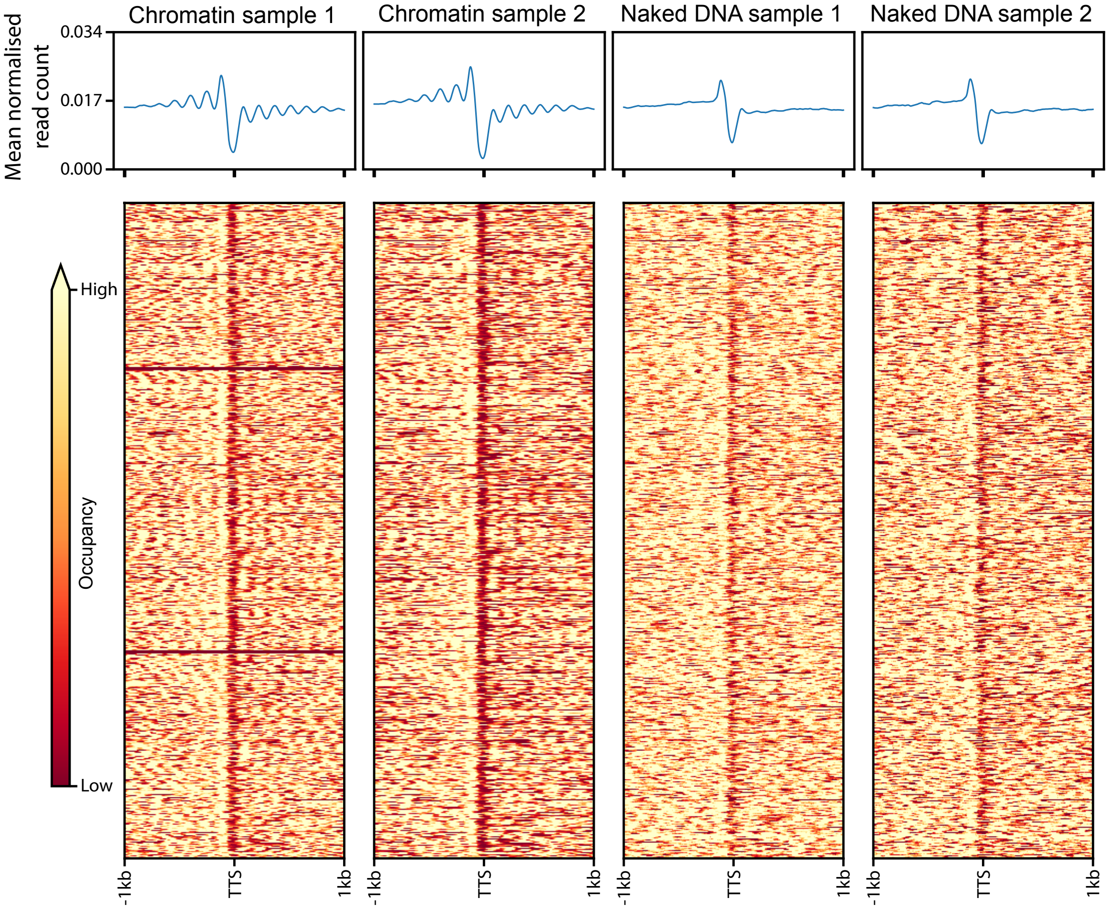

Mean normalised read count
0.034
0.017
0.000
Chromatin sample 1
Chromatin sample 2
Naked DNA sample 1
Naked DNA sample 2
High
Occupancy
Low
-1kb
TTS
1kb


**E**

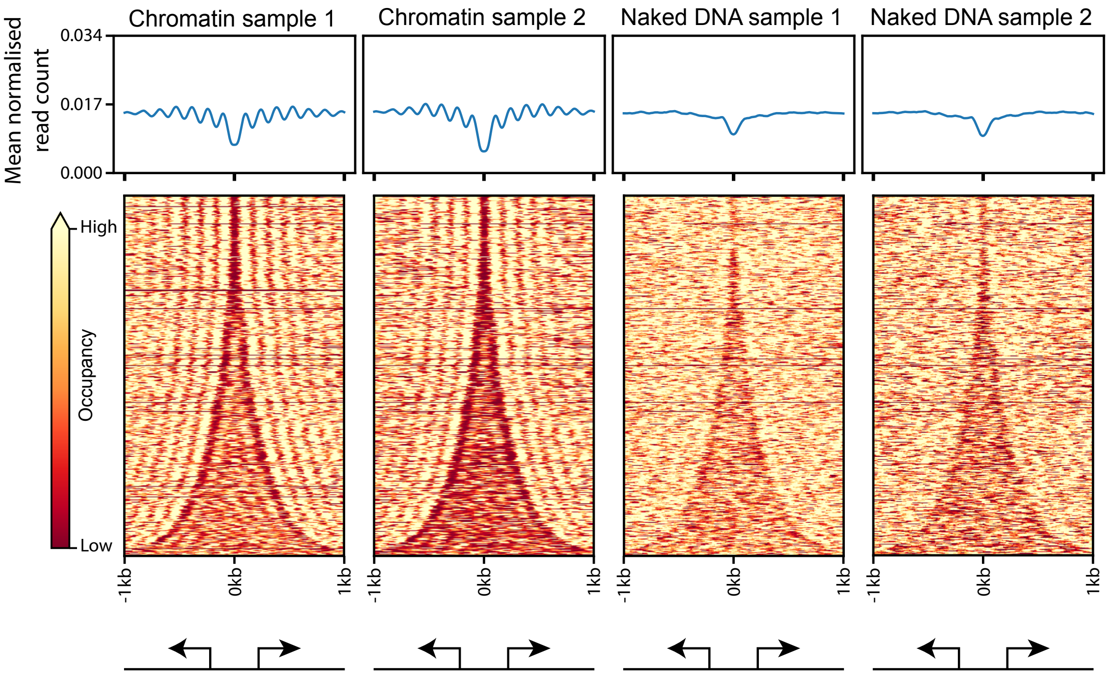

Mean normalised read count
0.034
0.017
0.000
Chromatin sample 1
Chromatin sample 2
Naked DNA sample 1
Naked DNA sample 2
High
Occupancy
Low
-1kb
0kb
1kb

F

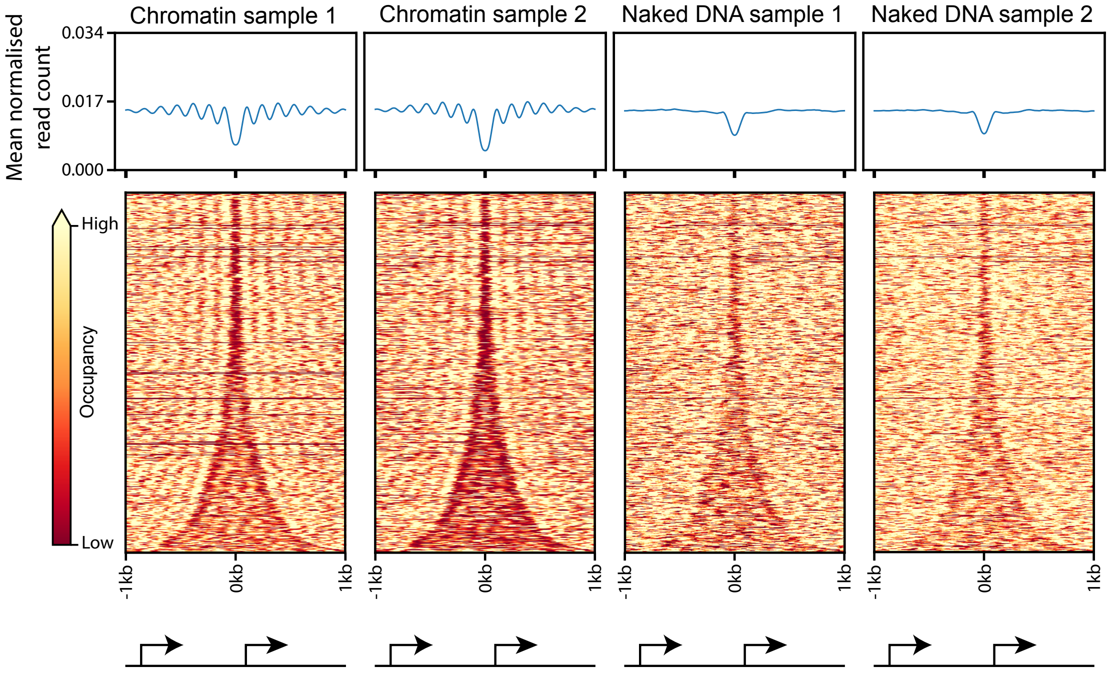
Chromatin sample 1
Chromatin sample 2
Naked DNA sample 1
Naked DNA sample 2
Mean normalised read count
0.034
0.017
0.000
High
Occupancy
Low
-1kb
0kb
1kb

G

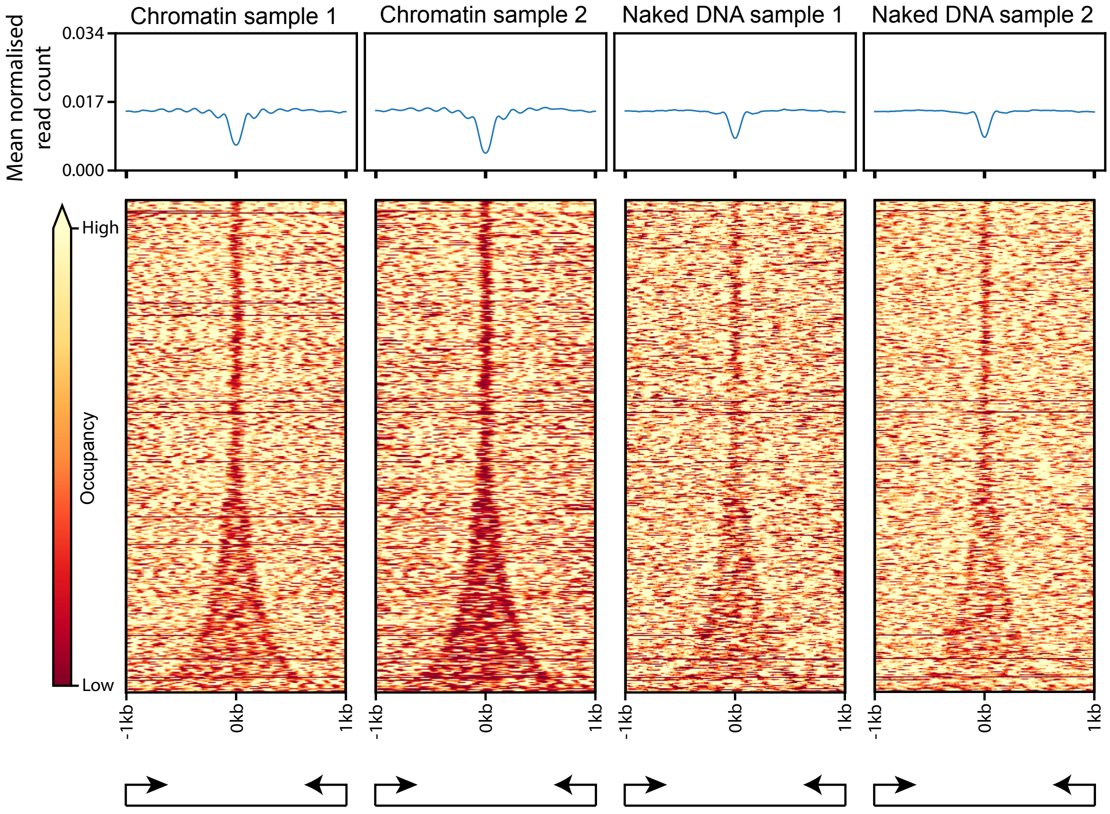
Chromatin sample 1
Chromatin sample 2
Naked DNA sample 1
Naked DNA sample 2
Mean normalised read count
0.034
0.017
0.000
High
Occupancy
Low
-1kb
0kb
1kb

H

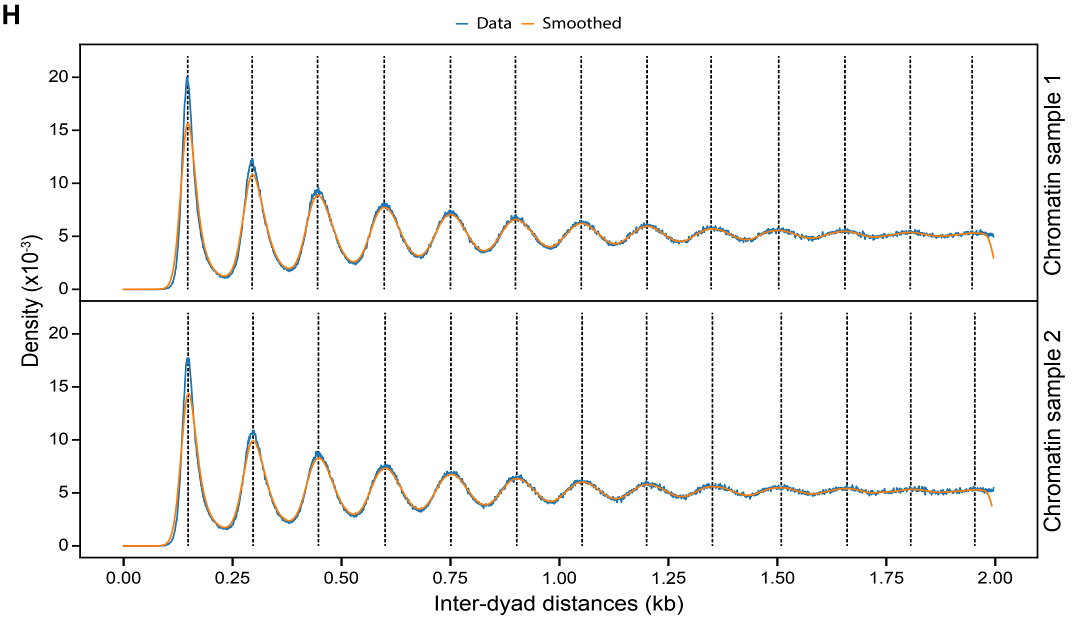

Data
Smoothed
Density (x10-3)
Chromatin sample 1
Chromatin sample 2
Inter-dyad distances (kb)
0.00
0.25
0.50
0.75
1.00
1.25
1.50
1.75
2.00
0
5
10
15
20


I

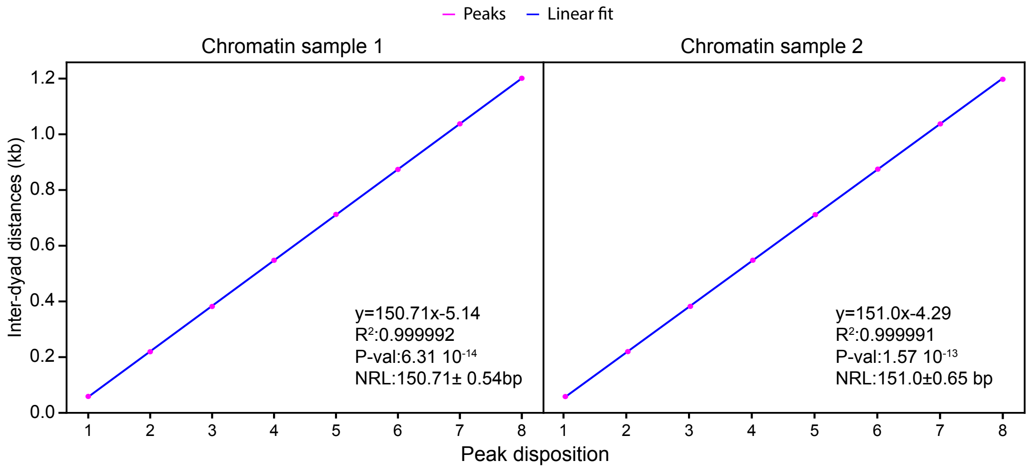

Peaks
Linear fit
Chromatin sample 1
Chromatin sample 2
Inter-dyad distances (kb)
Peak disposition
y=150.71x-5.14
R2:0.999992
P-val:6.31 10-14
NRL:150.71± 0.54bp
y=151.0x-4.29
R2:0.999991
P-val:1.57 10-13
NRL:151.0±0.65 bp


J

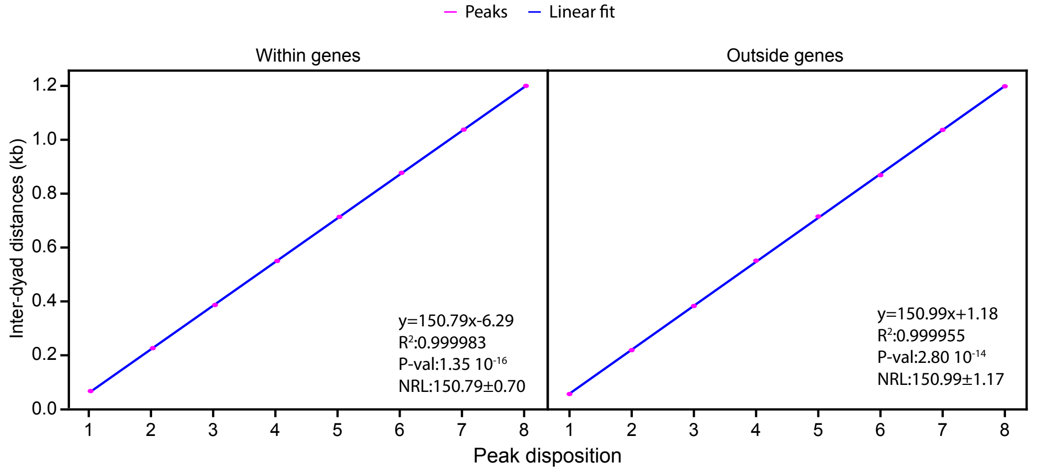

Peaks
Linear fit
Within genes
Outside genes
Inter-dyad distances (kb)
Peak disposition
y=150.79x-6.29
R2:0.999983
P-val:1.35 10-16
NRL:150.79±0.70
y=150.99x+1.18
R2:0.999955
P-val:2.80 10-14
NRL:150.99±1.17

**Figure S1. Nucleosome occupancy along the Mac *Paramecium* genome**

**(A)** MNase digestion of naked DNA with increasing MNase enzyme concentration. **(B)** Heatmap showing pairwise Pearson correlation between chromatin and naked DNA samples (two biological replicates for each). **(C)** Average profiles and heatmaps showing nucleosome occupancies around Transcription Start Sites (TSSs) in individual samples. **(D)** Average profiles and heatmaps showing nucleosome occupancies around Transcription Termination Sites (TTSs) in individual samples. **(E-G)** Average profiles and heatmaps showing nucleosome occupancies ±1 kb around the centre of intergenic regions. Pairs of genes have been divided into three groups based on their relative orientation: divergent (E), tandem (F) and convergent (G). Genes are sorted based on the intergenic distance. **(H)** Inter-centre distance between nucleosomes on the same scaffold for the two chromatin samples. In blue, distance distributions from actual data (from 1 bp to 2 kb, binning=1 bp); and in orange, the gaussian smoothed signal. Dashed black lines identify the local maxima (peaks centres) of the smoothed data (Materials and Methods). **(I)** In pink, the local maxima from Fig. S1 H ordered by increasing distance; in blue, the linear fitted model. On the bottom right of each panel, the information about the linear fitting and the estimated NRL for both chromatin replicates. P-values are calculated using a two-sided Z-test. **(J)** The same analysis as in Fig. 1 G by separating into the nucleosome centres overlapping with gene bodies (left panel) or outside of them (right panel). As in Fig. 1 G, local maxima are ordered by increasing distance. The relative linear fitting model is reported in blue. The bottom right gives the information about the linear fitting and the estimated NRL (Mean±SD). P-value is calculated using a two-sided Z-test.

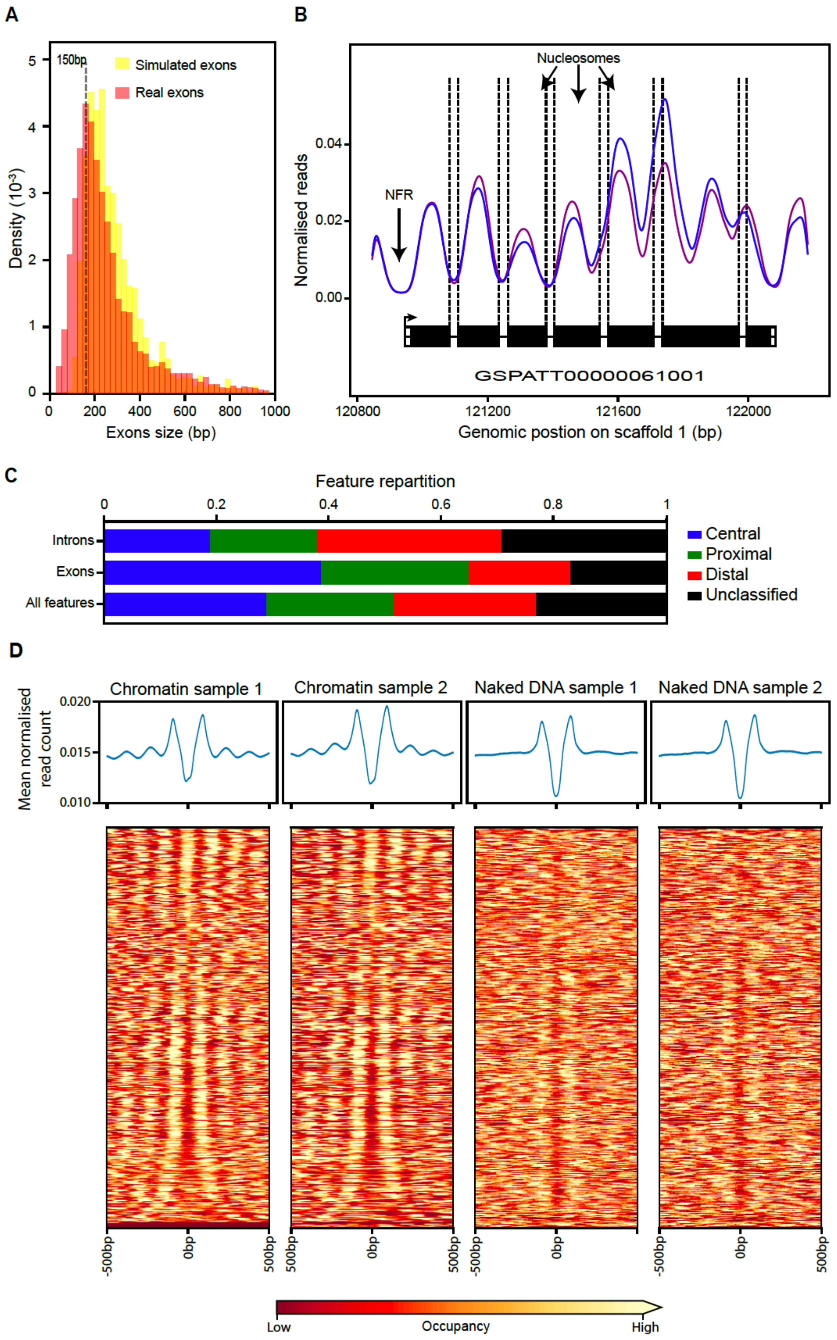


**Figure S2. Inter-nucleosomal DNA is frequently associated with intron position. (A)** Histogram showing exon size distribution (bin size = 25 bp) of transcription units whose extremities have been confirmed by 5' CAP-seq and poly A detection: in red, real exons; and in yellow, simulated exons created assuming uniform exon sizes within each transcript (see Materials and Methods) **(B)** The same genomic region as shown in Fig. 2 B. Tracks of two independent chromatin treated samples reporting nucleosome occupancy over genes with intron locations indicated by dashed vertical lines. Nucleosome free regions (NFRs) are found around the gene promoters and introns are frequently associated with inter-nucleosomal DNA. **(C)** Same figure as Fig. 2 G including features with distance >75 bp. "Unclassified" corresponds to introns and/or exons that do not belong to the three defined categories (central, proximal, distal). **(D)** Heatmap showing nucleosome occupancy ± 500 bp around intron centres in individual samples. Introns are ordered as in Fig. 2 D.

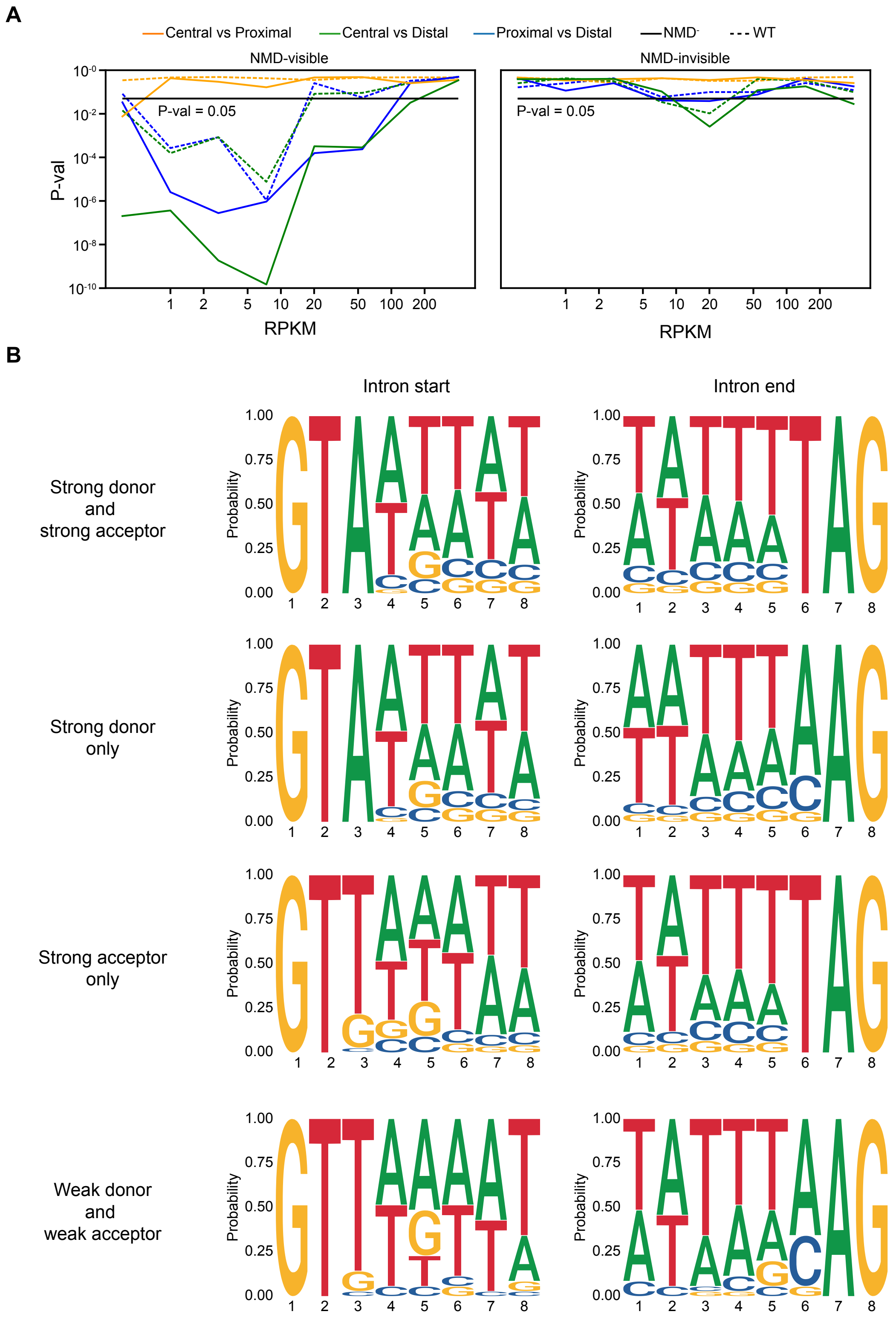


**Figure S3. Nucleosome positioning is associated with intron-splicing efficiency. (A)** P-values relative to Fig. 3 B were calculated using the Mann–Whitney U test and adjusted using false discovery rate. Comparisons of intron retention rates were evaluated in WT (dashed lines) and NMD-depleted cells (solid line), respectively, for NMD-visible introns (left) and NMD-invisible introns (right). The black line indicates P-value = 0.05**. (B)** Sequence logo plots showing the nucleotide composition of the splicing donors and acceptors sites relative to Fig. 3 C.

**A**

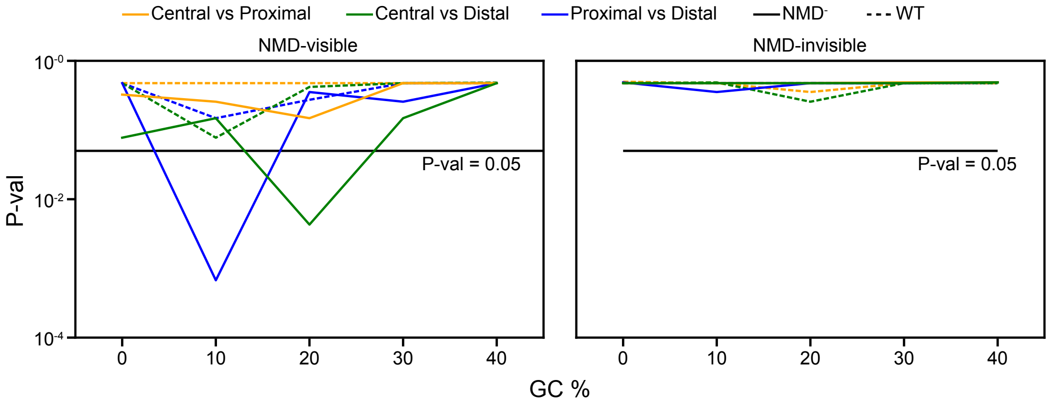


**B**

NMD-visible: $e^{SE}= 0.304 \log(T) - 0.165\, GC_I + 0.062\, P_I + 0.058\, N_I + 0.058\, L_I + 0.097\, L_T - 0.042\, L_{FE} + 0.023\, L_{PE} + 0.018\, D$

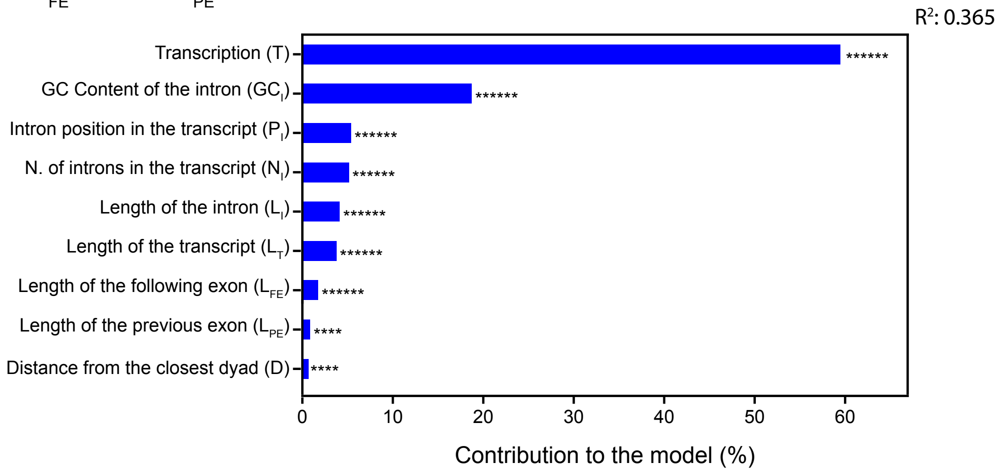


**C**

NMD-invisible: $e^{SE}= 0.295 \log(T) + 0.121\, L_T - 0.058\, GC_T - 0.051\, GC_I$

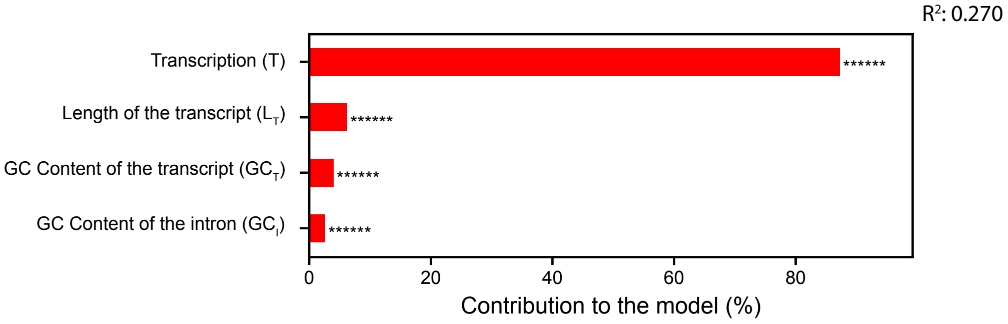


**Figure S4. GC content related to nucleosome positioning contributes to intron-splicing efficiency. (A)** P-values relative to Fig. 4 B were calculated using the Mann–Whitney U test and adjusted using false discovery rate. Comparisons of intron retention rates were evaluated in WT (dashed lines) and NMD-depleted (solid line) cells, respectively. Left: NMD-visible introns. Right: NMD-invisible introns. The black line indicates the position of P-value = 0.05**. (B-C)** Modelling Splicing Efficiency (SE) in NMD-depleted cells for the NMD-visible (B) and NMD-invisible (C) introns. The barplot reports the contribution to the model of each parameter. (P-value *<0.05, **< $10^{-2}$, ***< $10^{-3}$, ****< $10^{-4}$, *****< $10^{-5}$, ******< $10^{-6}$).
